# Disrupted stemness and redox homeostasis in mesenchymal stem cells of neonates from mothers with obesity: implications for increased adiposity

**DOI:** 10.1101/2025.04.14.648714

**Authors:** Sofía Bellalta, Erika Pinheiro-Machado, Jelmer Prins, Torsten Plösch, Paola Casanello, Marijke Faas

**Affiliations:** Department of Pathology and Medical Biology, University of Groningen, University Medical Center Groningen, Hanzeplein 1, EA11, 9713 GZ, Groningen, The Netherlands; Department of Obstetrics, School of Medicine, Pontificia Universidad Católica de Chile, Marcoleta 391, 8330024, Santiago, Chile; Department of Obstetrics and Gynaecology, University of Groningen, University Medical Center Groningen, Hanzeplein 1, 9713 GZ Groningen, The Netherlands; Perinatal Neurobiology Research Group, School of Medicine and Health Sciences, Carl von Ossietzky University of Oldenburg, Ammerländer Heerstraße 138, 26129, Oldenburg, Germany; Department of Neonatology, School of Medicine, Pontificia Universidad Católica de Chile, Marcoleta 391, 8330024, Santiago, Chile

**Keywords:** maternal obesity, stem cells, fetal programming, stemness properties, redox balance

## Abstract

Maternal obesity is a risk factor for increased fetal adiposity. The underlying mechanisms remain unclear, however, emerging evidence suggests that mesenchymal stem cells (MSCs), which are the precursors of adipocytes, from neonates of mothers with obesity exhibit enhanced adipogenic differentiation potential. We hypothesise that the MSCs of neonates from mothers with obesity have different stemness potential and redox state compared to the MSCs from mothers with normal weight. MSCs were isolated from neonates of women with obesity (BMI>30 kg/m², OB-MSCs) and women with normal weight (BMI <25 kg/m², NW-MSCs). OB-MSCs showed reduced stemness potential, as seen by a lower OCT3/4 expression and lower clonogenic capacity, than NW-MSCs (p<0.05). In addition, OB-MSCs showed higher levels of mitochondrial superoxide (O_2_^•-^), together with lower antioxidant SOD2 gene expression, compared to NW-MSCs (p<0.05). Conversely, OB-MSCs had higher levels of glutathione (GSH) compared to NW-MSCs (p<0.05). Upon exposure to H_2_O_2_ (250 μM), OB-MSCs displayed attenuated antioxidant response, with lower SOD1, SOD2 and GPX1 gene expression as compared to NW-MSCs (p<0.05). Upon exposure to higher oxidative stress (H_2_O_2_, 400 μM), total ROS levels were lower in OB-MSCs than in NW-MSCs. In contrast, when challenged for mitochondrial ROS, OB-MSCs showed higher levels of mitochondrial superoxide production as compared to NW-MSCs (p<0.05). Our results indicate that OB-MSCs have lower stemness potential, elevated mitochondrial O_2_^•-^ and a different basal and oxidative stress-induced redox profile compared to NW-MSCs. These changes in OB-MSCs could predispose them to an increase adipogeneic commitment.

**GRAPHICAL ABSTRACT:** 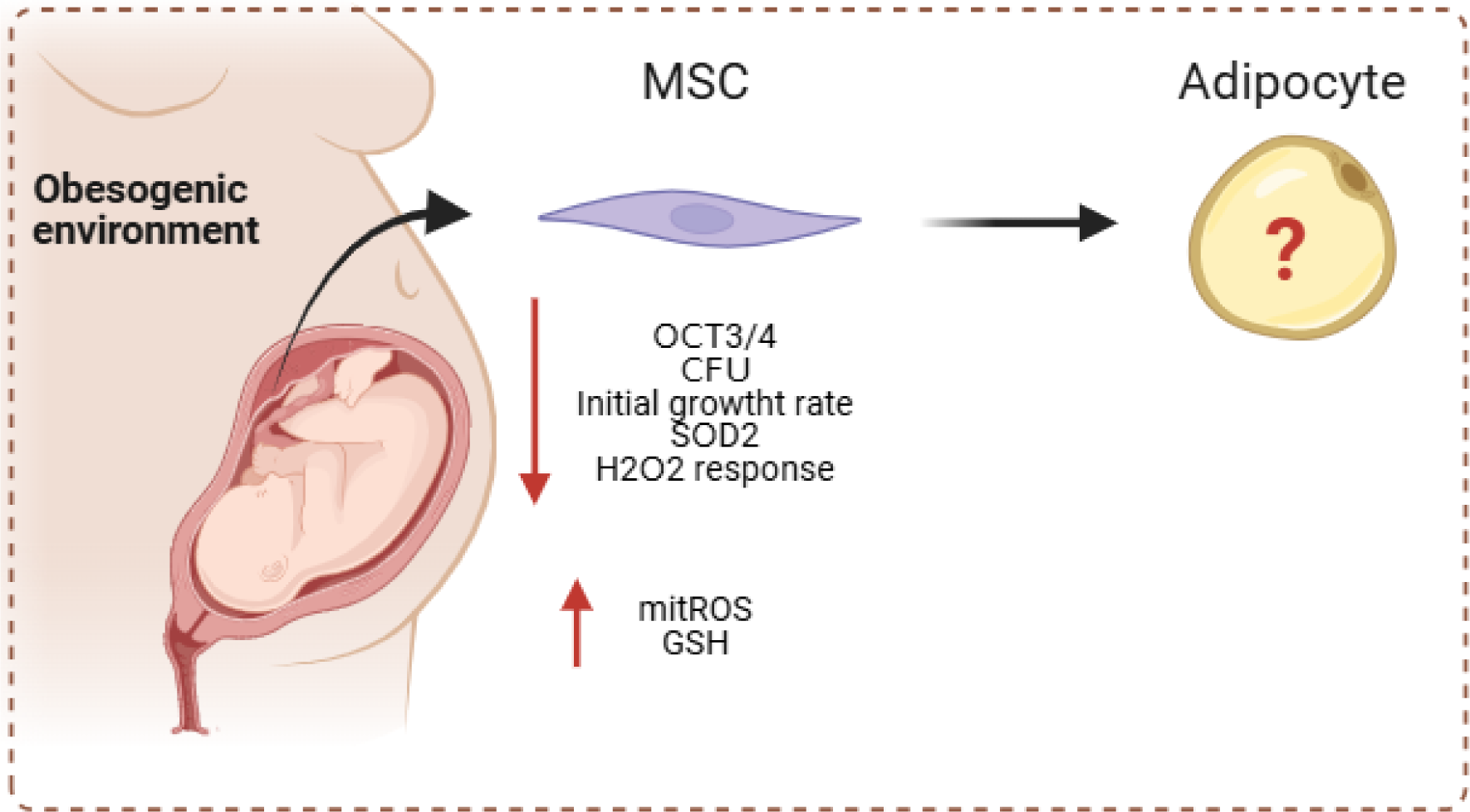

## INTRODUCTION

Maternal obesity leads to increased fetal growth and adiposity in the offspring [1–5]. The mechanisms that program fetal adiposity are still unclear; however, recent studies propose that mesenchymal stem cells (MSCs), the precursors of adipocytes, of neonates from mothers with obesity exhibit an enhanced potential for adipogenic differentiation [6–8]. This suggests that maternal obesity impacts MSCs, thereby influencing fetal adipogenesis.

MSCs are multipotent cells that give rise to the mesodermal cell lineages such as adipocytes, osteocytes, chondrocytes, and myocytes [9]. MSCs are established during early embryonic development and can be found in fetal tissues such as the Wharton’s jelly in the umbilical cord [10–13]. MSCs exhibit fundamental stem cell attributes: they are (1) capable of self-renewal and (2) retain several features reminiscent of embryonic stem cells [12, 14]. Together with (3) multipotency, they are collectively referred to as stemness properties [9, 12, 14]. These properties are influenced by the cellular environment, and may determine their future differentiation potential [15, 16].

Reactive Oxygen Species (ROS) are oxidants derived from molecular oxygen (O_2_) [17]. They are by-products of cellular aerobic metabolism but can also arise under pathological conditions [18]. ROS include the superoxide radical anion (O_2_^•-^) and other oxygen radicals, such as hydroxyl radical (·OH) and hydroxyl ion (OH^-^), but also non-radical derivatives such as hydrogen peroxide (H₂O₂) [17–19]. The primary source of intracellular ROS is the mitochondrial respiratory chain, among other cytoplasmatic sources [17, 20]. The redox balance –maintained through a dynamic equilibrium between ROS production and detoxification systems – is essential for managing oxidative stress [18, 19, 21].

Intracellular detoxification systems regulate ROS levels [19]. They include low-molecular-weight antioxidants such as glutathione (GSH) [22], together with antioxidant enzymes: superoxide dismutase (SOD) – responsible for modulating O_2_^•-^ levels – along with glutathione peroxidase (GPX) and catalase (CAT) – which regulate H₂O₂ levels [23]. Tight regulation of ROS is crucial for fundamental cellular processes such as proliferation and differentiation [18], particularly during adipogenic differentiation of MSCs. A physiological increase of ROS levels plays a critical role in driving MSC commitment to the adipogenic lineage [24–31, 32].

MSCs of neonates from mothers with obesity exhibit enhanced adipogenic differentiation potential. In this study, we hypothesise that maternal obesity also impacts stemness properties and redox state in MSCs, potentially contributing to their adipogenic commitment. To explore this, we evaluated stem cell multipotency, self-renewal capacity, proliferation, ROS levels and antioxidant mechanisms in neonatal MSCs from women with and without obesity.

## METHODS

### Study design

MSCs were isolated from the Wharton’s jelly of umbilical cords obtained from neonates of mothers with normal weight and mothers with obesity. These MSCs were analysed to assess their fundamental characteristics and redox parameters. We used MSCs in basal state to measure stemness by evaluating differentiation potential and expression of Stage Specific Embryonic Antigen 4 (SSEA4), Octamer binding protein 3/4 (OCT3/4), clonogenicity and proliferation. We also measured ROS and O ^•-^ levels in MSCs derived from both study groups. To investigate antioxidant mechanisms, we focused on basal GSH levels and gene expression levels of antioxidant enzymes SOD1/2, GPX1 and CAT in MSCs. Additionally, MSCs were subjected to incubation with 250 μM H₂O₂ to evaluate the response of antioxidant enzymes to oxidative stress. MSCs were also incubated with 400 μM H₂O₂, 100 μM tert-butyl hydroperoxide (tBHP) or 20 μM Antimycin A, to evaluate intracellular ROS or O ^•-^ production in response to oxidative stress (Supplementary Figure 1).

### Subjects

Umbilical cords were anonymously obtained from the placentas of women with obesity or normal weight, who delivered at the University Medical Centrum Groningen (UMCG) maternity ward, Groningen, The Netherlands, from August 2021 to February 2023. Umbilical cords were obtained from biological waste material following routine childbirth procedures. No specific ethical permission was required as tissue was classified as medical waste and was de-identified, with no donor information collected. The study complied with all relevant institutional and national ethical regulations regarding the use of biological materials. All procedures were also conducted according to the Helsinki Declaration. A pregestational maternal body mass index (BMI) >30 kg/m² was considered for the group of women with obesity (OB, n=15) and 18.5-24.5 kg/m² for women with normal weight (NW, n=15). The inclusion criteria included women >18 years, single and term pregnancies (>37 weeks). The exclusion criteria included women with gestational diabetes, preeclampsia, preterm birth, and neonatal complications. All characteristics of the mother and newborn were blinded, apart from the woman’s pregestational BMI (weight, height) and gestational weight.

### Isolation of Wharton’s jelly-derived MSCs

Umbilical cords were obtained and immediately processed to acquire MSCs primary cultures. MSCs were isolated using the explant method [12, 33]. Therefore, the umbilical cord was washed in cold Dulbecco’s Phosphate Buffered Saline (DPBS, Gibco, Thermo Fischer Scientific) and cut into three cm pieces inside a laminar flow cabinet. Each piece was cut longitudinally, and blood vessels were discarded. Wharton’s jelly explants were plated and cultured with Dulbecco’s modified Eagle medium containing 10 mM glucose (DMEM, Gibco, Thermo Fischer Scientific), 10% Fetal calf serum (FCS, Sigma-Aldrich), 5.000 UI/mL Penicillin-Streptomycin (Gibco, Thermo Fischer Scientific), and maintained at 37°C in 5% CO₂. Media was changed every four days, and culture was maintained for 10-14 days. By this time, a solid population of cells had sprouted out from the explants and had covered the explant perimeter. Subsequently, the sprouted cells were trypsinized (Gibco, Thermo Fischer Scientific) and expanded into further passages by seeding at 4.000/cm² in culture flasks. The medium was changed every three days, and cells were maintained in the same culturing conditions. Cells were passaged after six days of culture (passages 1-3). Unless otherwise stated, all experiments were performed in fresh cells on passages 2-3.

### Immunophenotyping

Surface markers were characterized according to the International Society of Cell Therapy criteria [34]. Briefly, cells were detached with trypsin and centrifuged at 2500 x g for three minutes. After this, 1×10^6^ cells were resuspended in 100 μl of FACS buffer (2% FCS in DPBS) in FACS tubes (Falcon, Corning) and incubated for 30 minutes in the dark with antibodies for CD73 (APC, 1:50, Miltenyi Biotec), CD90 (PE, 1:50, Miltenyi Biotec), CD105 (BV421, 1:20, BioLegend), CD34 (PE-Dazzle 594, 1:20, Biolegend), CD45 (BV785, 1:20, BioLegend), and CD11b (APC-Vio770, 1:50, Miltenyi Biotec). Then, cells were washed with 2 ml FACS buffer and centrifuged at 2250 x g for three minutes, twice. Finally, cells were resuspended in 300 μL FACS buffer. Samples were measured by Novocyte Quanteon II Flow Cytometer (Agilent), and data was analysed in NovoExpress (Agilent). Events were gated for side scatter height (SSC-H) vs forward scatter height (FSC-H) to exclude debris and forward scatter height (FSC-H) vs forward scatter area (FSC-A) to exclude doublets. Further, populations were gated in the fully stained samples, using gate setting in the unstained samples.

### Lineage commitment

Cells were seeded in a 24-well plate at 10.000 cells/cm². When cells were 80% confluent, they were incubated with differentiation media for each lineage using the following induction media: a) adipogenic induction medium: DMEM (25 mM glucose; Gibco, Thermo Fischer Scientific) supplemented with 10% FCS (Sigma-Aldrich), 1 nM insulin (Lonza Bioscience), 0.5 mM 3-isobutyl-1-methylxanthine (Sigma-Aldrich), 0.1 μM dexamethasone (Sigma-Aldrich); b) osteogenic induction medium: DMEM (25 mM glucose, Gibco, Thermo Fischer Scientific) supplemented with 10% FCS (Sigma-Aldrich), 10 mM β-glycerophosphate (Sigma-Aldrich), 0.1 μM dexamethasone (Sigma-Aldrich), 0.05 mM ascorbic acid (Sigma-Aldrich); and c) myogenic induction medium: DMEM (25 mM glucose, Gibco, Thermo Fischer Scientific) supplemented with 10% Fetal calf serum (FCS, Sigma-Aldrich), 0.1% TGF-β1 (PeproTech). The medium was changed every three days, and differentiation was carried out through 14 (osteogenic/myogenic/adipogenic) and 21 days (adipogenic). After this time, cells were fixed in the plate with 2% PFA (Sigma-Aldrich). Experiments were performed in duplicates.

### Staining of lineage commitment

For adipocyte staining, fixed cells in the plate were washed with water, rinsed with 60% 2-propanol, and immediately stained with Oil red O (Sigma-Aldrich) for 30 minutes. Nuclei were briefly stained with hematoxylin (Sigma-Aldrich). For osteoblasts staining, fixed cells were washed with DPBS and stained in 0.5% Alizarin Red (Sigma-Aldrich) for ten minutes. Nuclei were stained with hematoxylin. Images were acquired by Leica DM IL Led microscope (Leica Microsystems). For smooth muscle staining, fixed cells were washed with DPBS and incubated with Phalloidin-AlexaFluor488 (1:250; Thermo Fischer Scientific) and DAPI (1:5000, Roche Diagnostics) for 30 minutes. Images were acquired with EVOS FL (Life Technologies, Fischer Scientific).

### SSEA-4 labelling

Cells in passage 2 were seeded at 2.000 cells/cm² in Lab-Tek chambers (Thermo Fischer Scientific) and fixed with 2% PFA (Sigma-Aldrich). After fixation, cells were washed with DPBS (Gibco, Thermo Fischer Scientific) and incubated with SSEA-4 (1:500, Invitrogen) for two hours. After this, cells were washed and incubated with a secondary anti-mouse Alexa Fluor 546 (Thermo Fischer Scientific) antibody for one hour. Cells were further stained with DAPI (Roche Diagnostics) for 15 minutes. Samples were washed with DPBS, and images were taken by EVOS FL (Life Technologies, Fischer Scientific). Fluorescence intensity was quantified by Image J (NIH) considering: [Fluorescent intensity = mean cell intensity – mean background intensity]. Background fluorescence was measured considering the area of each cell [35].

### RNA isolation

Cells were cultured in 6-well plates at 5.000 cells/cm². For the induced state, we used an oxidative challenge with 250 μM H₂O₂ (Merck) for six hours. Pilot experiments indicated that this challenge induced antioxidant gene expression without affecting cell viability (Supplementary Figure S2). After reaching confluency, cell lysates were collected with TRIzol (Invitrogen). Total RNA was isolated as described by the manufacturer [36]. Briefly, the sample was homogenized, and layers were separated with chloroform (Merck). RNA was precipitated from the aqueous layer with 2-propanol (Merck). The pellet was reconstituted in water, and RNA concentration and purity were measured with a NanoDrop ND-100 UV-Vis spectrophotometer (NanoDrop Technologies).

### cDNA synthesis

A total of 500 ng of total RNA was used for reverse transcriptase cDNA synthesis. The reverse transcription (RT) was performed in total RNA following the standard protocol for RevertAid Reverse Transcriptase (Thermo Fischer Scientific Inc.). Briefly, samples were treated with DNAse I (Thermo Fischer Scientific Inc.) and incubated with Random Hexamer Primers, dNTPs set, Ribolock RNAse Inhibitor and RevertAid Reverse Transcriptase (Thermo Fischer Scientific) in a thermocycler (Biometra, Analytik Jena) following standard protocol.

### OCT3/4 and antioxidant enzymes gene expression

Quantitative PCR (qPCR) was performed with TaqMan Assay for OCT 3/4 (POU5F1 Hs04260367_gH, Thermo Fischer Scientific) according to the manufacturer’s instructions [37]. Glyceraldehyde 3-phosphate dehydrogenase (GAPDH Hs02786624_g1, Thermo Fischer Scientific) was used as the housekeeping gene. For antioxidant enzymes, qPCR was performed with FastStart Universal SYBR Green Master (Roche Diagnostics) for SOD1, SOD2, GPX1 and CAT according to manufacturer’s instructions. β-2-Microglobulin (B2M), Glyceraldehyde 3-phosphate dehydrogenase (GAPDH) and 60S ribosomal protein P0 (RPLP0) were used as housekeeping genes. qPCR was performed in triplicates in Viia7 Real-time PCR System (Thermo Fischer Scientific). Data analysis was carried out using Viia7 Software (Thermo Fischer Scientific). The threshold cycle (Ct) for gene amplification was detected and normalized as a relative expression using 2^delta Ct (ΔCt = Ct [gene of interest] – Ct [average housekeeping genes]) (Supplementary table S1 – primers list).

### Colony Forming Unit assay

Cells were seeded in low density (10 and 100 cells/cm²) for passage 2 and passage 5. The media was changed every three days. After 14 days, cells were fixed in 2% PFA (Sigma-Aldrich) and stained with 0,05% Crystal Violet (Sigma-Aldrich). Images were acquired by Leica DM IL Led microscope (Leica Microsystems). Colonies with more than 50 cells were considered positive colonies. Experiments were performed in duplicates.

### Growth kinetics

MSCs were seeded on 12-well plates at 2.000 cells/cm². The medium was changed every three days for ten days. Every 48 hours, two arbitrary wells were trypsinized (Gibco, Thermo Fischer Scientific), and cells were counted in triplicate with a Coulter Counter Z2 (Beckman Coulter Diagnostics).

### Total intracellular ROS levels

Basal and induced total intracellular ROS levels were assessed with 2’,7’-dichlorodihydrofluorescein diacetate (H2DCFDA) (Thermo Fischer Scientific) [38]. Briefly, cells were seeded at 3.000/cm² in 24-well plates and cultured until confluence. After reaching confluency, cells were detached with trypsin (Gibco, Thermo Fischer Scientific) and centrifuged at 1200 x g for five minutes. After this, cells from each well were resuspended in 1 ml of DPBS (Gibco, Thermo Fischer Scientific) in FACS tubes (Falcon, Corning), and incubated with H2DCFDA (5 μM) for 30 minutes at 37°C in the dark. For the prooxidant challenge, cells were incubated for the last 15 minutes with 400 μM of hydrogen peroxide (H₂O₂, Merck) as a physiological ROS, or with 100 μM of tBHP (Sigma-Aldrich) as a stable non-physiological ROS, at 37°C in the dark. Untreated cells were incubated at the same time. Cells were washed with 1 ml DPBS, centrifuged at 1200 x g for five minutes and resuspended in 300 μL DPBS. Then, 0.1 μM of ZombieNIR (BioLegend) was added to evaluate cell viability. Five thousand events were measured in the FITC channel in the Novocyte Quanteon II Flow Cytometer (Agilent). Events were gated for SSC-H vs FSC-H, and FSC-H vs FSC-A to exclude doublets and dead cells, respectively (Supplementary Figure S3). Data was analysed with NovoExpress (Agilent).

### Total intracellular and mitochondrial O_2_ ^•-^ levels

O_2_^•-^ levels were evaluated with dihydroethidium (DHE) and MitoSOX (Thermo Fischer Scientific) [38]. Briefly, cells were seeded at 3.000/cm² for each condition in 24-well plates and cultured until confluence. After reaching confluency, cells were detached with trypsin (Gibco, Thermo Fischer Scientific) and centrifuged at 1200 x g for five minutes. After this, cells were resuspended in 1 ml of DMEM (Gibco, Thermo Fischer Scientific) without FBS, in FACS tubes (Falcon, Corning) and incubated with DHE (5 μM) or mitoSOX (1 μM) for 30 minutes at 37°C in the dark. For the pro-oxidative challenge, cells were incubated in suspension for the last 15 minutes with 10 μM of Antimycin A (Sigma-Aldrich)at 37°C in the dark. Untreated cells, i.e. without pro-oxidative challenge, were incubated at the same time. For each condition, cells were washed with 1 ml DPBS, centrifuged at 1200 x g for five minutes and resuspended in 300 μL DPBS. Then, 0.1 μM of ZombieNIR (BioLegend) was added for cell viability assessment. Five thousand events were measured in the PE channel by Novocyte Quanteon II Flow Cytometer (Agilent). Events were gated for SSC-H vs FSC-H, and FSC-H vs FSC-A to exclude doublets, and dead cells were excluded. (Supplementary Figure S3). Data was analysed with NovoExpress (Agilent).

### Glutathione levels

Cells were seeded at 5.000 cells/cm² in 6-well plates and cultured in standard conditions until confluence. After reaching confluency, cells were detached with trypsin (Gibco, Thermo Fischer Scientific) and centrifuged at 900 x g for five minutes. 1x 10^5 cells were resuspended in a buffer containing 50 mM sodium phosphate and 1 mM EDTA, sonicated and stored at -80°C. After collecting all samples, they were thawed and deproteinized with 5% meta-phosphoric acid (Sigma-Aldrich). Oxidized (GSSG), reduced (GSH), and total glutathione (GSHt) levels were measured with a Glutathione GSH/GSSG Assay kit (Sigma-Aldrich), following the manufacturer’s instructions. The optical density of samples was measured in a 96-well plate reader (Biotek Epoch 2, Agilent) at 412 nm wavelength (time zero and ten minutes). Data was calculated according to the standard curve. For GSSG, the process was performed in the presence of the scavenger 1-methyl-2-vinylpyridinium triflate. Finally, total GSH levels were calculated with the following formula:

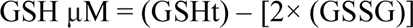

### Statistical analysis

The parameters investigated were expressed as the median value and 25th and 75th percentiles in Graphpad Prism (Graphpad Inc). Mann–Whitney U tests were used to compare between NW-MSCs and OB-MSCs. Data was log transformed, and a two-way ANOVA test was used to assess independent variables BMI category (NW-MSCs or OB-MSCs) and passaging for clonogenic capacity assays (Passages 2 and 5), mRNA expression, ROS or O ^•-^ levels, followed by a Tukey’s range test. P values < 0.05 were considered statistically significant.

## RESULTS

### Wharton’s jelly-derived cells express MSCs markers and have multipotent potential

Wharton’s Jelly tissue explants from all subjects showed outgrowth of adherent MSCs (data not shown). Cells from both group were >88% positive for MSC markers CD73 and CD90, and CD105; and negative for hematopoietic/leukocyte markers CD11b, CD34 and CD45 (Figure 1A). There was no difference in the expression of any marker between NW-MSCs and OB-MSCs (Table 1). Both NW-MSCs and OB-MSCs exhibited calcium deposit accumulation upon osteogenic differentiation (Alizarin Red staining) and displayed alpha-smooth muscle actin fibers (α-SMA) after myogenic differentiation (Phalloidin staining), and were positive for lipid accumulation (Oil red O staining) when induced with adipogenic induction (Figure 1B).

**Figure 1.**
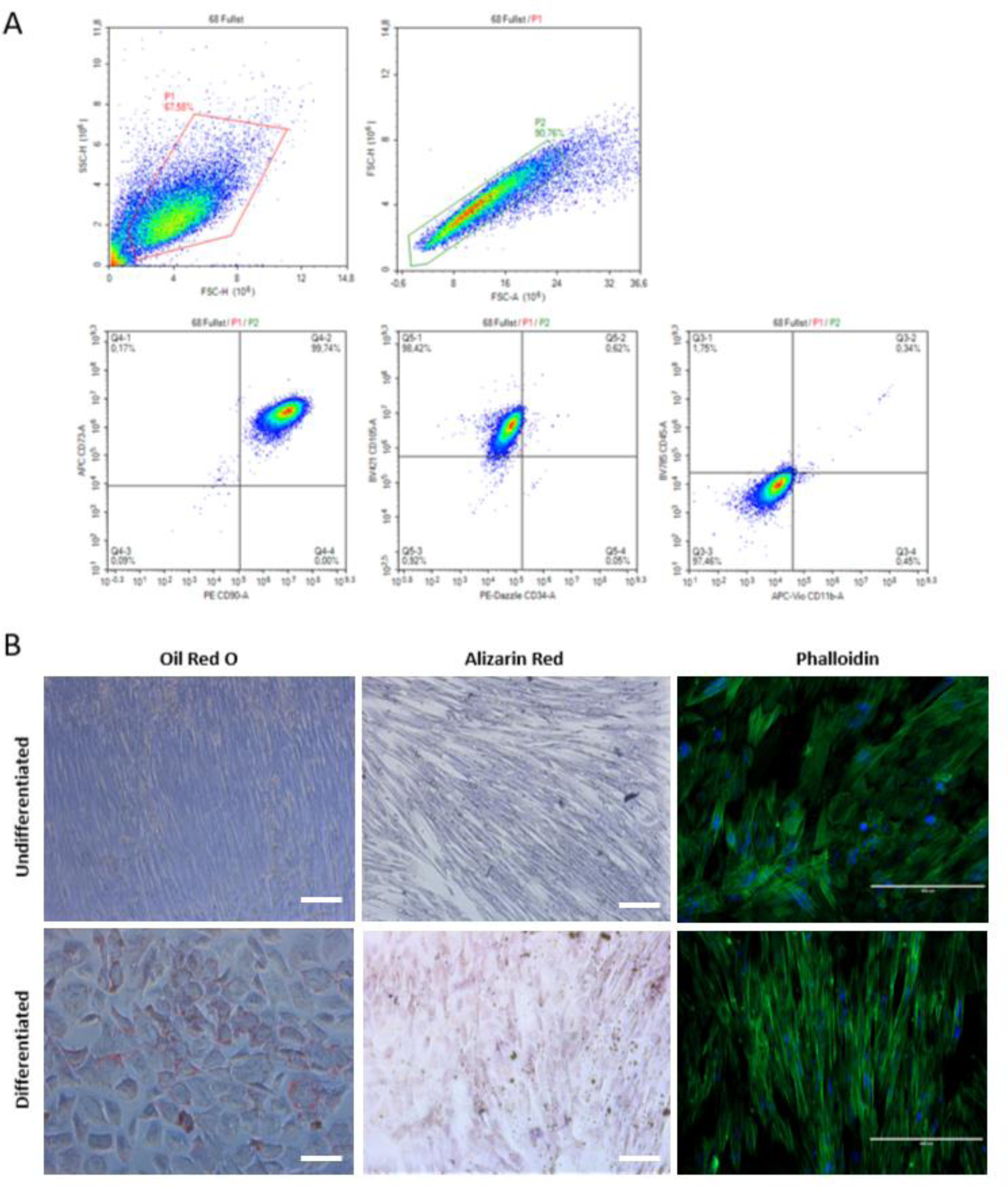
Wharton’s jelly-derived MSCs exhibit MSCs characteristics. **A**. Cells were positive for surface markers CD73, CD90, and CD105, while negative to CD34, CD45 and CD11b in NW-MSCs and OB-MSCs (n=3). The gating of cells is plotted as forward scatter *versus* side scatter for cell population and forward scatter versus forward scatter for singlets (top). Cell populations CD73+ and CD90+, CD105+ and CD34-, and CD45- and CD11b-(bottom). Representative graphs show surface markers of NW-MSCs. **B.** Cells were cultured in three different induction media (osteogenic and myogenic) for 14 days, and in adipogenic induction medium for 21 days. Cells were stained for lipids (Oil Red O), calcium (Alizarin Red) and α-SMA (phalloidin) (n=3). Representative images show induction of NW-MSCs (Scale bars Oil Red O and Alizarin Red: 100 μm; Phalloidin: 400 μm).

**Table 1.** Wharton’s jelly-derived MSCs exhibit MSCs immunophenotype.

| **Antigen** | **NW MSCs (%)** | **OB MSCs (%)** | **P value** |
| --- | --- | --- | --- |
| CD73 | 99.75 ± 0.1 | 99.77 ± 0.2 | 0.99 |
| CD90 | 97.53 ± 2.8 | 97.73 ± 3.0 | 0.83 |
| CD105 | 88.20 ± 4.0 | 89.42 ± 3.8 | 0.50 |
| CD34 | 0.18 ± 0.03 | 0.21 ± 0.1 | 0.99 |
| CD45 | 0.45 ± 0.3 | 0.65 ± 0.2 | 0.56 |
| CD11b | 0.28 ± 0.1 | 0.99 ± 0.7 | 0.30 |

P value of Mann-Whitney U test; n=3.

### OB-MSCs exhibit reduced stemness markers and properties compared to NW-MSCs

NW-MSCs and OB-MSCs were positive for SSEA4. There was no difference in the fluorescence intensity between both groups (Figure 2A). However, gene expression of OCT3/4 was decreased in OB-MSCs compared to NW-MSCs (p=0.03) (Figure 2B). NW-MSCs showed 15.3 ± 8.6 colonies, compared with 14.3 ± 3.5 colonies in OB-MSCs for P2. In P5, NW-MSCs exhibited 7.6 ± 1.5, compared with 2.8 ± 0.7 colonies in OB-MSCs. There was an effect of the passage over the number of colonies (p<0.0001), with a decrease in the number of colonies in passage 2 compared to 5 only for OB-MSCs (p=0.002, Figure 2C).

**Figure 2.**
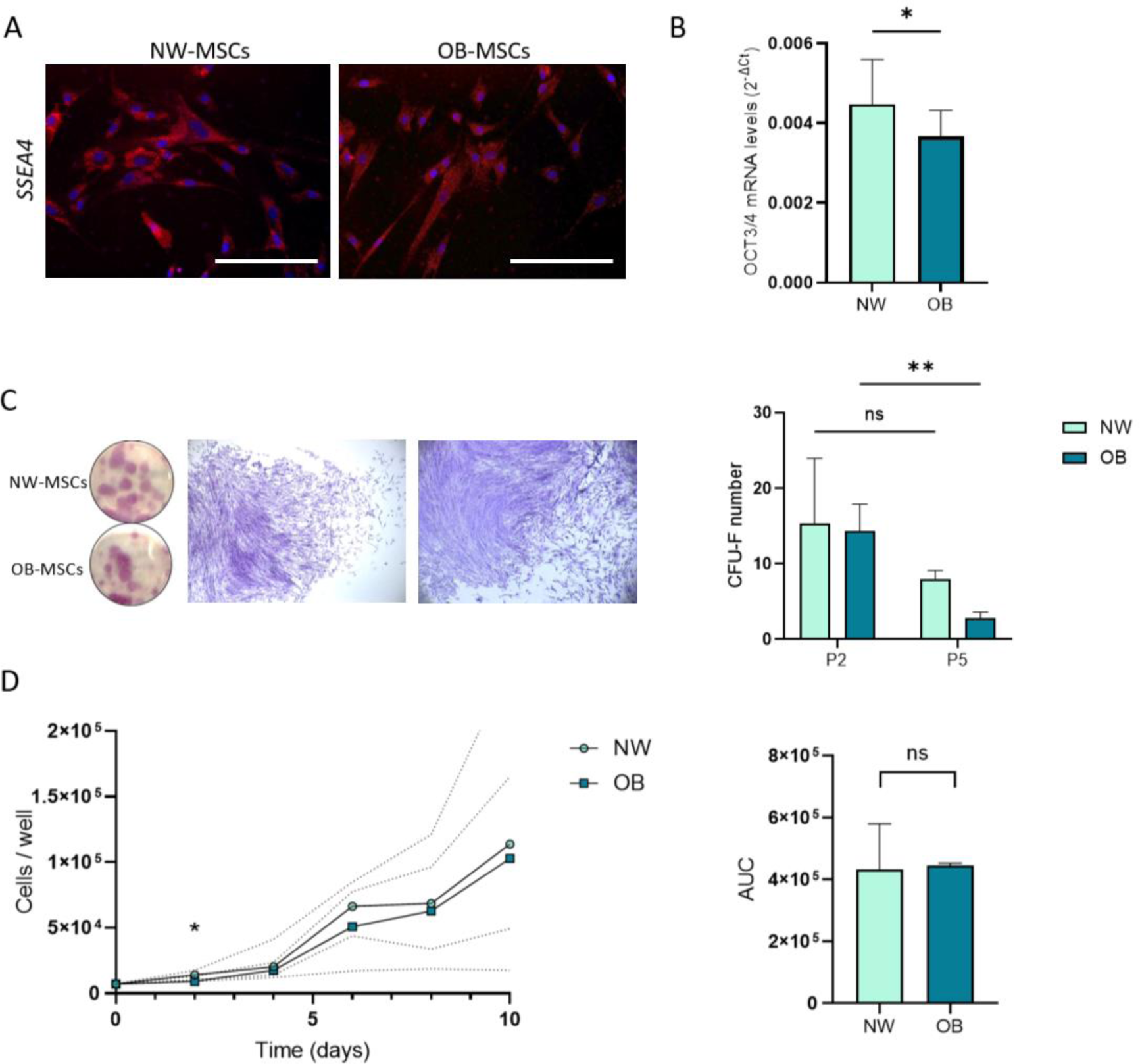
OB-MSCs exhibit lower stemness properties, clonogenic capacity and growth rate. **A.** Representative images of MSCs from normal weight and obesity groups stained for Stage Specific Embryonic Antigen 4 (SSEA-4) (scale bar: 100 μM). **B.** OB-MSCs showed lower gene expression levels of the stemness marker OCT3/4, compared to NW-MSCs (n=5). **C**. Representative images of colonies per well with magnification (left) and quantification of positive colonies at day 14 (right, n=6). **D.** Cells were seeded at the same starting density and counted every 2 days. Growth curve was calculated considering cells per well each day (left, n=5). Area under the curve (AUC) of NW-MSCs and OB-MSCs for 10 days (right). *p< 0.05 Mann-Whitney U test; **p< 0.01 Tukey’s range post hoc test.

### OB-MSCs have a lower initial growth rate compared to NW-MSCs

NW-MSCs and OB-MSCs proliferated for ten days from 7.000 to 116.333 ± 49.511 cells and 94.583 ± 70.732 cells, respectively. The area under the curve in NW-MSCs was 425.400 ± 140.396, whereas it was 307.000 ± 194.664 in OB-MSCs. This difference was not significant (Figure 2D). However, OB-MSCs proliferated to 9.300 ± 2.167 cells, whereas NW-MSCs exhibited significantly higher number of cells (13.733 ± 2.644 cells) on day 2 (p =0.01). Further, on days 4, 6, 8 and 10, there was no difference in the number of cells between the two groups (Figure 2D).

### OB-MSCs show higher mitochondrial oxidative stress compared to NW-MSCs in basal state

To evaluate basal ROS levels, we measured total intracellular ROS, total intracellular O_2_ ^•-^ and mitochondrial O_2_ ^•-^ in NW-MSCs and OB-MSCs (Figure 3). Total intracellular ROS and total intracellular O_2_ ^•-^ levels were not different between both groups (p=0.39 and p=0.1). However, there was an increase in the levels of mitochondrial O_2_ ^•-^ of OB-MSCs compared to NW-MSCs in the basal state (p = 0.03, Figure 3).

**Figure 3.**
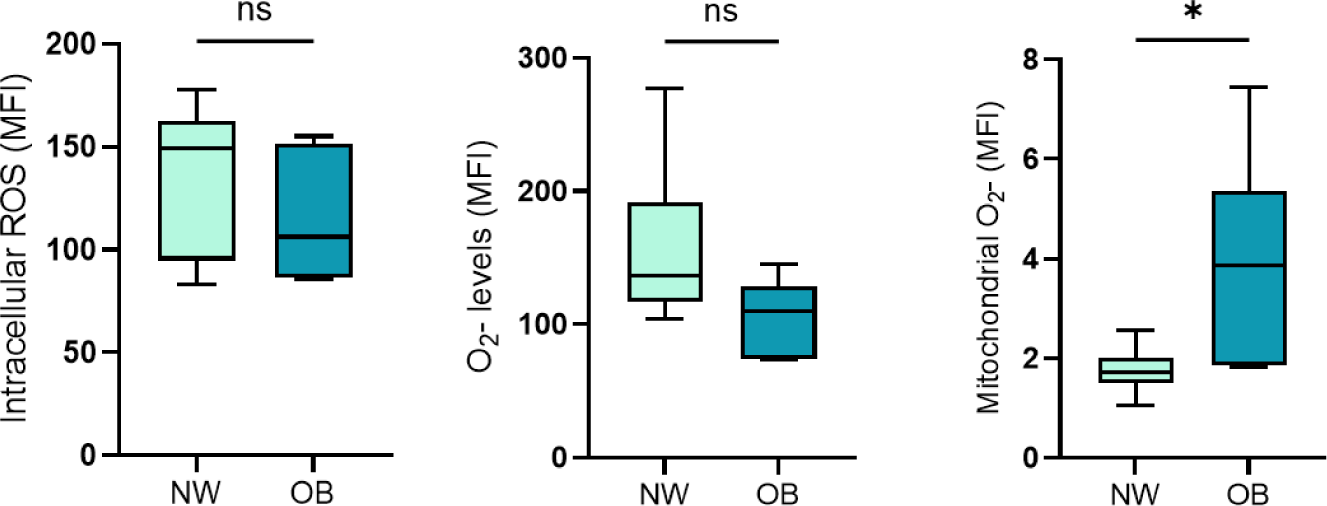
Basal redox state in NW-MSCs and OB-MSCs. Basal ROS levels: cells were incubated with H2DCFDA for total intracellular ROS, DHE for total O_2_^•-^ or mitoSOX for mitochondrial O_2_^•-^, for 30 minutes.. * p<0.05 (Mann-Whitney U test, median ± range, n=6)

### Basal antioxidant gene expression is decreased in OB-MSCs compared to NW-MSCs

We evaluated the mRNA expression of the genes of antioxidant enzymes in the basal state. OB-MSCs showed lower basal mRNA expression of SOD2 as compared with NW-MSCs (p=0.03), with no differences in SOD1, GPX1 and CAT between OB-MSCs and NW-MSCs (p=0.2, 0.1 and 0.1, respectively, Figure 4A).

**Figure 4.**
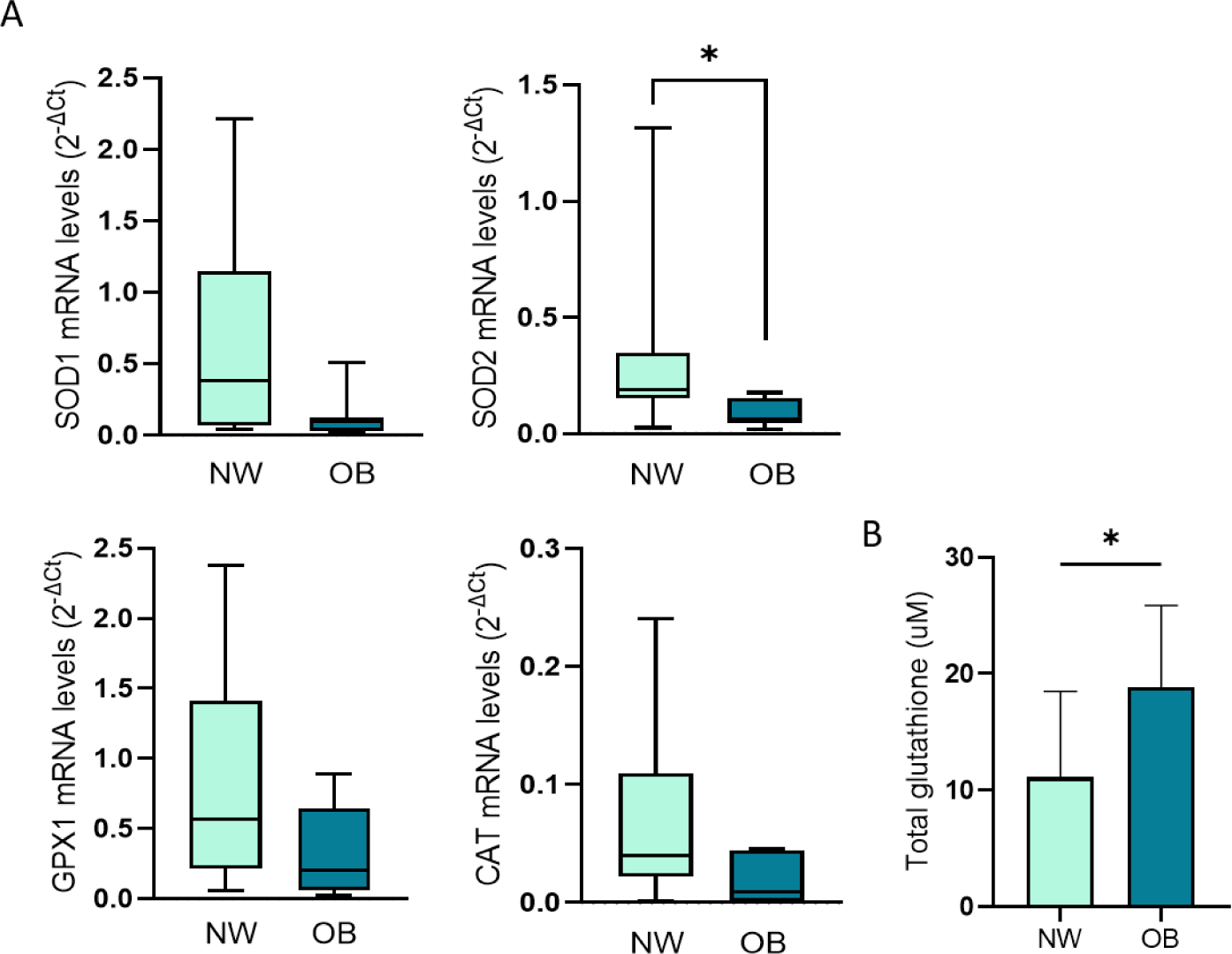
Basal antioxidant mechanisms in NW-MSCs and OB-MSCs. **A.** SOD1/2, GPX1 and CAT gene expression were measured in both groups. **B.** Glutathione intracellular levels in NW-MSCs and OB-MSCs. * p<0.05 (Mann-Whitney U test, median ± range, n=7).

### OB-MSCs show higher antioxidant GSH levels compared to NW-MSCs in basal state

Levels of GSH were 20.4 ± 4.9 μM in OB-MSCs, and 12.6 ± 3.61 μM in NW-MSC, indicating that there are higher basal glutathione levels in OB-MSCs, suggesting a higher redox buffering capacity (Figure 4B). We did not find differences in the ratio of GSH/GSSG between NW-MSCs and OB-MSCs (data not shown), which is another indicator of the glutathione buffering capacity.

### OB-MSCs, compared to NW-MSCs, exhibit lower levels of intracellular ROS in response to a pro-oxidative challenge

To evaluate the intracellular ROS response and overall antioxidant response, we performed a challenge with H₂O₂ (400 μM) and tBHP (100 μM) (Figure 5A). There was an effect of H₂O₂ (p=0.0001) and tBHP (p<0.0001) challenges, and of maternal obesity (p=0.01). Moreover, there was a significant increase of intracellular ROS after treatment with H₂O₂ compared to untreated in NW-MSCs, while this was not seen for OB-MSCs (p=0.001 and p=0.1, respectively). Additionally, results show an effect of tBHP treatment compared to untreated cells for both NW-MSCs and OB-MSCs (p=0.0004 and 0.01 in NW-MSCs and OB-MSCs, respectively). Intracellular ROS levels in response to the H₂O₂ challenge were lower in OB-MSCs compared to NW-MSCs (p=0.03, Figure 5A).

**Figure 5.**
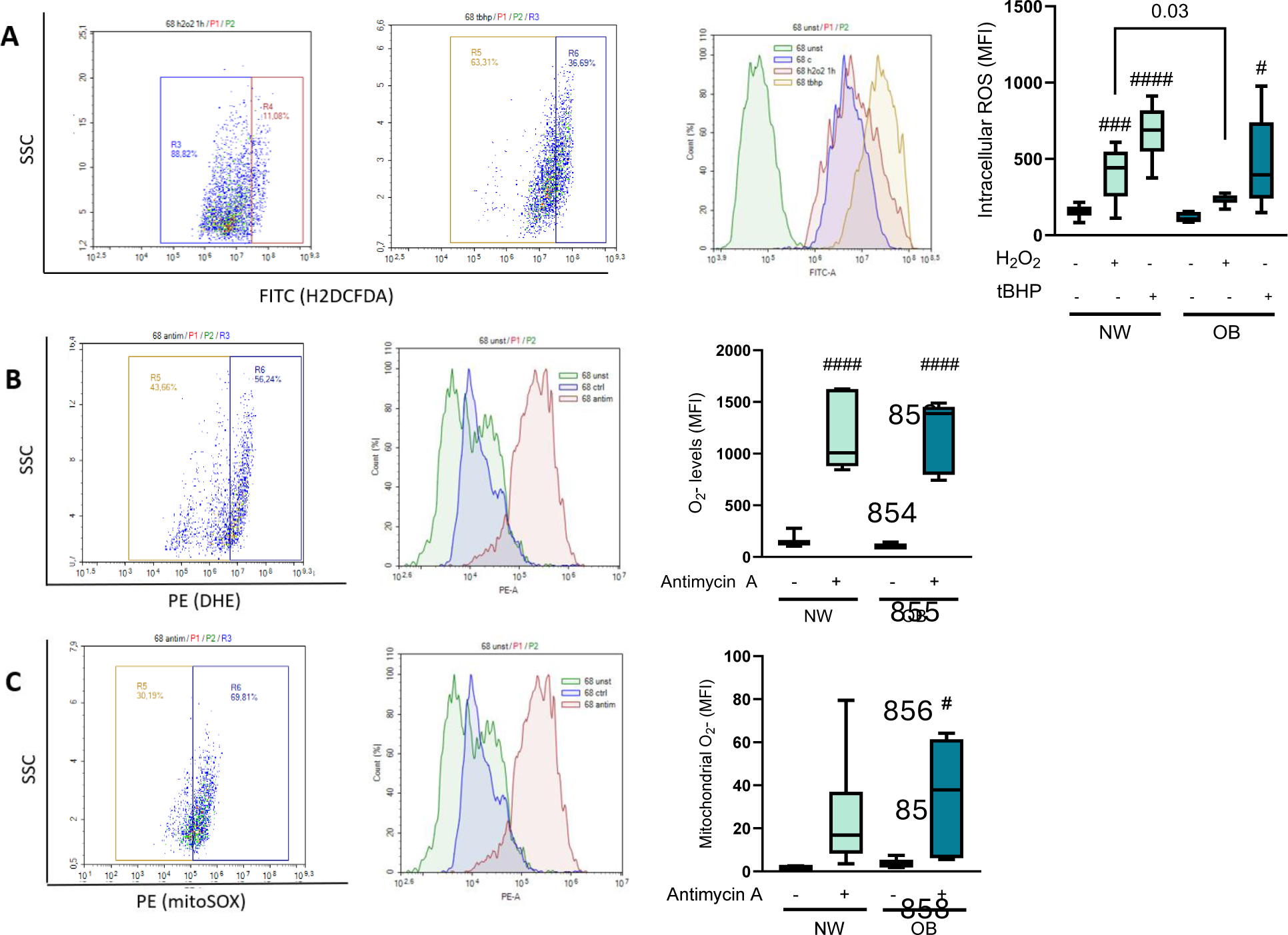
Induced ROS levels in NW-MSCs and OB-MSCs. **A.** Cells were incubated with H2DCFDA after incubation with or without H₂O₂ (400 μM) or tBHP (100 μM) for 15 minutes to assess induced ROS levels. Two-way ANOVA: Effect of H₂O₂ (p=0.0001), tBHP (p<0.0001) and obesity (p=0.01) (median ± range, n=6). **B**. Cells were incubated with DHE after incubation with or without Antimycin A (10 μM) for 15 minutes to assess O_2_^•-^. Two-way ANOVA: Effect of Antimycin A (p<0.0001) and obesity (p=0.9) (median ± range, n=6). **C.** Cells were incubated with mitoSOX after incubation with or without Antimycin A (10 μM) for 15 minutes to assess mitochondrial O_2_^•-^. Two-way ANOVA: Effect of Antimycin A (p=0.002) and obesity (p=0.4) (median ± range, n=6). For all graphs, Tukey’s range post hoc test comparisons vs basal condition: # p<0.05; ###p<0.001; ####p<0.0001).

### NW-MSCs and OB-MSCs increase intracellular O ^•-^ production in response to a pro-oxidant mitochondrial challenge

When challenged with antimycin A, there was an increase in the total intracellular O_2_ ^•-^ levels in both groups (Figure 5B). The results show an effect of antimycin A treatment (p<0.0001), but not of maternal obesity (p=0.7). Additionally, there was a significantly increased O_2_ ^•-^ after treatment, compared to untreated cells for both groups (p<0.0001 NW-MSCs and OB-MSCs, Figure 5B).

### NW-MSCs and OB-MSCs increase mitochondrial O_2_ ^•-^ levels production in response to a pro-oxidative mitochondrial challenge

For mitochondrial O_2_^•-^, there was an effect of antimycin A (p=0.002) but not of maternal obesity (p=0.4, Figure 5C). Furthermore, results show that there was an increase in O_2_^•-^ levels in OB-MSCs in antimycin A treatment compared with no treatment (p=0.04), while there was no difference in NW–MSCs in treated versus non-treated cells (p=0.1, Figure 5C).

### Induced antioxidant gene expression is decreased in OB-MSCs compared to NW-MSCs

In response to a challenge with H₂O₂, there was an overall effect of maternal obesity over antioxidant gene expression levels (SOD1: p=0.001; SOD2: p<0.0001; GPX1: p<0.0001; CAT: 0.003). Further, SOD1, SOD2 and GPX1 mRNA significantly increased compared to the basal, unstimulated state in NW-MSCs (p=0.04, 0.02 and 0.03, respectively). None of the enzyme’s mRNA increased after the H₂O₂ challenge compared to the basal state in OB-MSCs (Figure 6). The CAT gene expression was unaffected by the H₂O₂ challenge in both groups.

**Figure 6.**
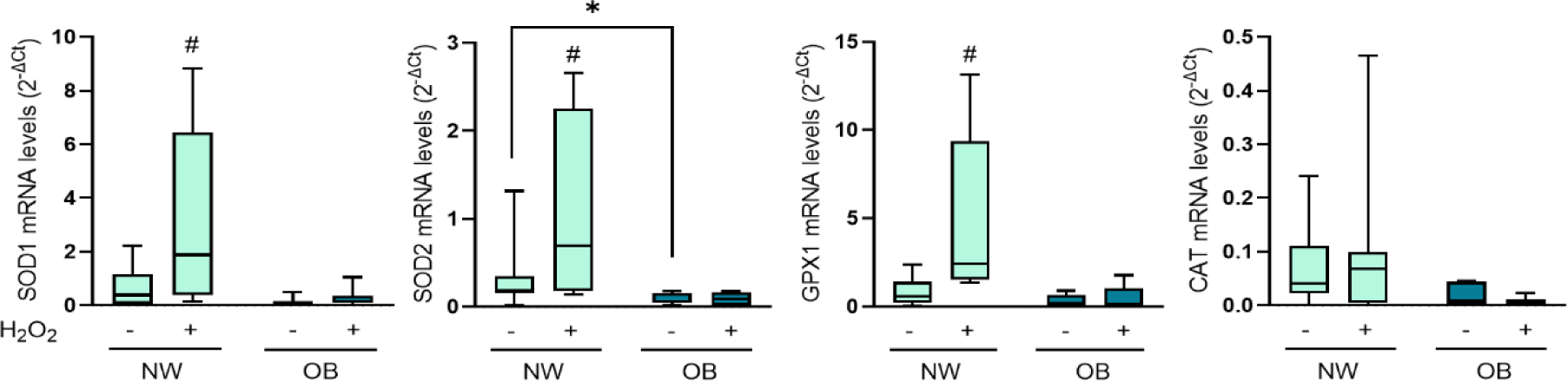
Induced antioxidant gene expression in NW-MSCs and OB-MSCs. SOD1/2, GPX1 and CAT gene expression were measured for basal and H₂O₂-induced conditions (250 μM) (median ± range, n=7). Two-way ANOVA: Effect of BMI (SOD1: p=0.001; SOD2: p<0.0001; GPX1: p<0.0001; CAT: p=0.003. *p< 0.05 Tukey’s range post hoc test NW-MSCs vs OB-MSCs. #p< 0.05 Tukeýs range post hoc test basal vs induced

## DISCUSSION

The primary objective of this study was to evaluate stemness signatures and redox balance in adipocyte progenitor cells, the MSCs, from neonates of women with normal weight and obesity. We found that OB-MSCs have a decreased self-renewal, clonogenic capacity and initial growth rate compared to NW-MSCs. In terms of the basal redox status, the OB-MSCs show higher mitochondrial ROS, and a decreased expression of SOD2 compared to NW-MSC. Regarding the redox capacity after a pro-oxidative challenge, OB-MSCs exhibited a decreased response to an oxidative challenge in terms of antioxidant enzyme gene expression. Interestingly, OB-MSCs showed an increased in GSH levels, paralleled by lower levels of intracellular ROS after H2O2 exposure, compared to NW-MSCs.

In our study, we found lower expression of the pluripotency marker OCT3/4 in OB-MSCs compared with NW-MSCs. This indicates a diminished self-renewal potential compared to NW-MSCs [39, 40] This suggests that OB-MSCs decrease their pluripotency and self-renewal capacity, and was translated into a reduced clonogenic capacity in OB-MSCs upon passaging. The significant reduction in clonogenic capacity, coupled with their reduced initial proliferative ability, are characteristics commonly associated with an aging phenotype of stem cells [41–43]. Prior studies have highlighted the role of aging and cellular senescence signaling in impairing adipogenesis and contributing to features of metabolic dysfunction [7, 44, 45]. Thus, we propose that OB-MSCs may exhibit characteristics of cellular aging, and future studies should further investigate aging- and senescence-related markers, together with their potential involvement in the increased adipogenic potential of OB-MSCs.

We found that OB-MSCs exhibit higher mitochondrial O ^•-^ levels compared to NW-MSCs, indicating mitochondrial stress. These data align with findings of Iaffaldano *et al*., who reported lower oxygen consumption rates and a reduced mitochondrial count in OB-MSCs compared to NW-MSCs [8]. A lower oxygen consumption rate is known to increase electron leakage under various energetic conditions, which, in turn, leads to elevated mitochondrial O_2_^•-^ production [46, 47]. Furthermore, Baker *et al*. characterized mitochondrial gene expression in OB-MSCs and noted significant changes during adipogenic differentiation *in vitro*. Their study highlighted upregulation of the mitochondrial respiratory chain, alongside downregulation of mitochondrial biogenesis and mitophagy in OB-MSC versus NW-MSCs [48]. Another study reported distinct epigenetic regulation of mitochondrial gene expression in OB-MSCs [49]. Collectively, our findings show mitochondrial dysfunction and mitochondrial oxidative stress in OB-MSCs.

SOD2 is downregulated in OB-MSCs compared to NW-MSCs in the basal state, suggesting decreased basal antioxidant capacity. SOD2, a manganese-dependent superoxide dismutase (MnSOD), is the main antioxidant enzyme that scavenges O ^•-^ in the inner mitochondrial matrix and acts as a first line of defence against mitochondrial oxidative damage [23]. The reduced basal levels of SOD2 are in line with the elevated mitochondrial O ^•-^ observed in OB-MSCs, indicating mitochondrial dysfunction in these cells. Furthermore, inhibition of complex III by antimycin A, considered as a mitochondrial stressor, led to higher O ^•-^ levels in OB-MSCs compared to the response of NW-MSCs, which also suggests the presence of mitochondrial stress in OB-MSCs. Altered mitochondrial function plays an important role in the pathophysiology of obesity-induced disorders, therefore, our findings provide evidence that mitochondrial dysfunction is already present in neonatal progenitor cells, which may contribute to the development of metabolic diseases later in life [50–53].

Our findings reveal elevated levels of total GSH in OB-MSCs compared to NW-MSCs. We propose that the increase in GSH may be a compensatory response to the blunted activity of GPX1. When GPX1 activity is diminished, GSH consumption decreases, leading to its accumulation [54]. This buildup of GSH could serve as a protective mechanism to counter oxidative stress despite impaired GPX1 function [55–57]. Higher levels of GSH are consistent with reports in individuals with obesity [56, 58, 59]. Thus, elevated synthesis of GSH may serve as a compensatory mechanism for regulating cytosolic ROS levels. Indeed, when exposed to 400 μM H₂O₂, OB-MSCs exhibited lower intracellular H₂O₂ levels compared to NW-MSCs. Coupled with the attenuated response of GPX1 the 250 μM H₂O₂ challenge in OB-MSCs compared to NW-MSCs, these findings suggest alterations in glutathione metabolism in OB-MSCs.

We evaluated general cytosolic antioxidant response by stimulating cells with prooxidant challenges: 400 μM of H₂O₂ and tBHP. Our results show that OB-MSCs have lower intracellular ROS levels in response to the H₂O₂ challenge, compared to NW-MSC, suggesting that primary antioxidant mechanisms are more efficient in OB-MSCs compared to NW-MSCs in response to this prooxidative challenge. We suggest that the higher levels of GSH in OB-MSCs may play a protective role during this H₂O₂ challenge. However, we do not discard other redox mechanisms that may participate in cellular detoxification, such as peroxiredoxins, thioredoxins and other low molecular mass scavengers [19], which should be considered for future studies. When the MSCs were subjected to a stronger prooxidant challenge, tBHP, which is more stable in aqueous solutions [60], no significant differences in intracellular ROS were observed between NW-MSCs and OB-MSCs. These findings suggest that the antioxidant response in OB-MSCs may be adapted to withstand milder oxidative challenges, such as H₂O₂ exposure. However, both groups lack the capacity to effectively counteract more potent oxidants like tBHP.

An increase in oxidant levels can induce adaptation to stress and enhance cellular resilience, a phenomenon known as hormesis or mitohormesis [61, 62]. This mechanism increases tolerance, thereby preventing excessive strain on the system while promoting the life span [17, 62]. This effect is particularly evidenced in early life stressors due to the plasticity of individuals [61–64]. In our study, OB-MSCs showed a blunted oxidative stress-response of antioxidant enzymes SOD1, SOD2 and GPX1 gene expression in OB-MSCs. Whether this blunted antioxidant response in OB-MSCs has an overall beneficial effect on tolerating ROS is a question that remains.

We thus demonstrate that maternal obesity can impact the stemness properties and redox balance in neonatal MSCs. These modifications may represent a mechanism by which maternal obesity impacts fetal progenitor cells and adipogenesis, contributing to the predisposition of the offspring to metabolic disorders later in life. The limitation of our study is that we could not collect clinical data from the patients. Future studies should integrate maternal and neonatal clinical parameters, which could provide valuable insights into donor variability and could link MSCs characteristics to obesity in the offspring. Future studies should also focus on exploring the specific molecular pathways behind redox balance and adipogenic potential to provide deeper understanding of the association between maternal obesity and offspring adiposity.

## ETHICS APPROVAL

Not applicable.

## CONSENT TO PARTICIPATE

Not applicable.

## DATA AVAILABILITY

Data will be made available on request.

## CODE AVAILABILITY

Not applicable.

## AUTHOR’S CONTRIBUTIONS

SB: Conceptualization, methodology, investigation, formal analysis, writing. EPM: Investigation, methodology. JP: Methodology. PC: Conceptualization. TP: Review and editing. MMF: Review and editing.

## SUPPLEMENTARY MATERIAL

**Supplementary Table S1.**
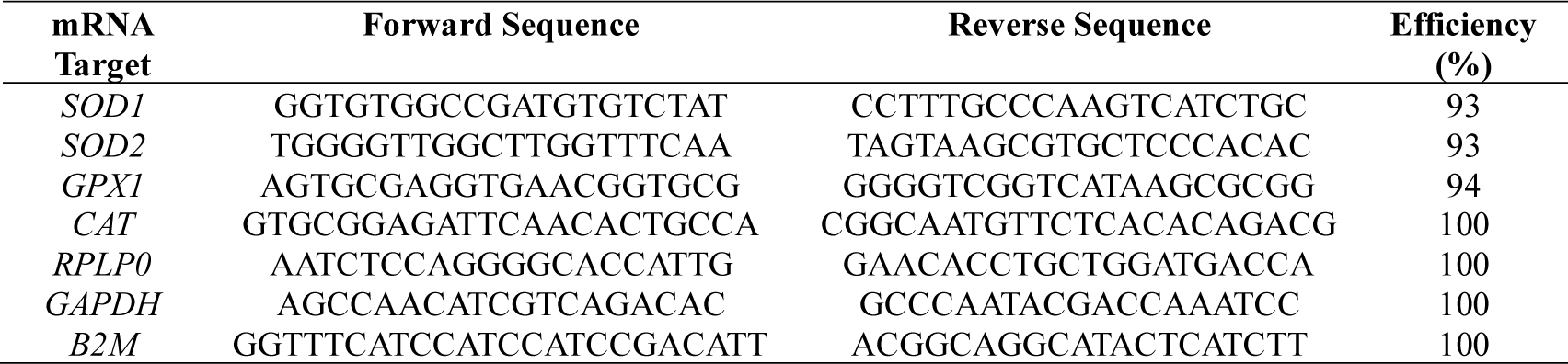
Primers sequences and amplification conditions for RT-qPCR.

**Supplementary Figure S1.**
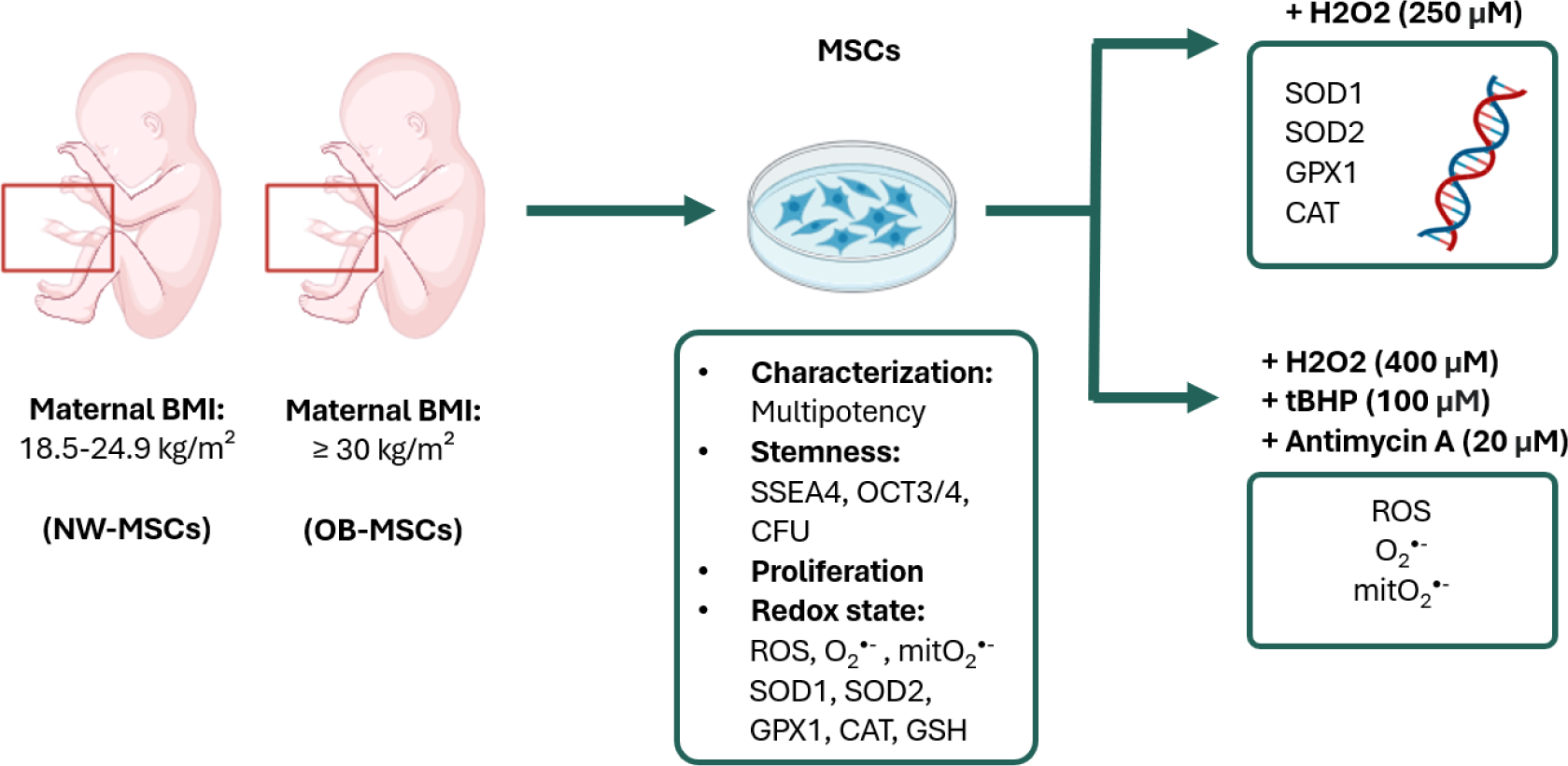
Study design. MSCs were isolated from the Wharton’s jelly of umbilical cords obtained from neonates of mothers with normal weight (NW-MSCs=15) and mothers with obesity (OB-MSCs=15). These MSCs were cultured characterized and further analysed to assess their stemness potential: SSEA4, OCT3/4, clonogenicity; and proliferation capacity. Further, we assessed redox parameters: reactive oxygen species (ROS), superoxide (O_2_^•-^), mitochondrial superoxide (mitO_2_^•-^) levels, gene expression of antioxidant enzymes and glutathione. Additionally, MSCs were subjected to pro-oxidative challenges: 250 μM H₂O₂ was used to induce changes in the gene expression of antioxidant enzymes, while 400 μM H₂O₂, 100 μM tert-butyl hydroperoxide (tBHP) or 20 μM Antimycin A, were used to induce intracellular ROS or O_2_^•-^.

**Supplementary Figure S2.**
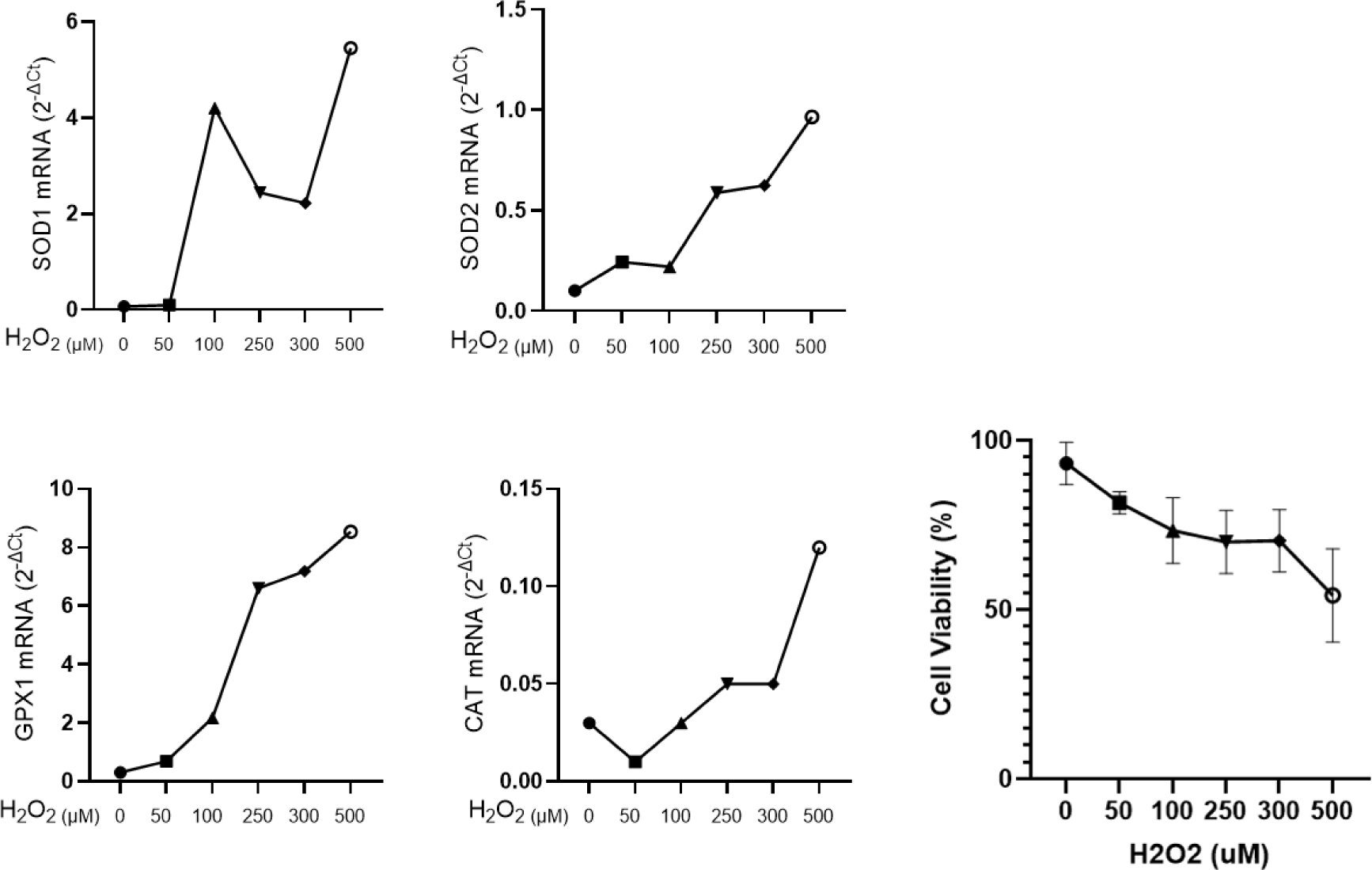
H₂O₂ challenge and viability assays in NW-MSCs. Concentration-response to H₂O₂ experiments in NW-MSCs for standardization of an oxidative challenge. Left: gene expression of SOD1, SOD2, GPX1 and CAT was assessed after six hours of incubation with H_2_O_2_. Right: cell viability was evaluated with trypan blue in response to H₂O₂.

**Supplementary Figure S3:**
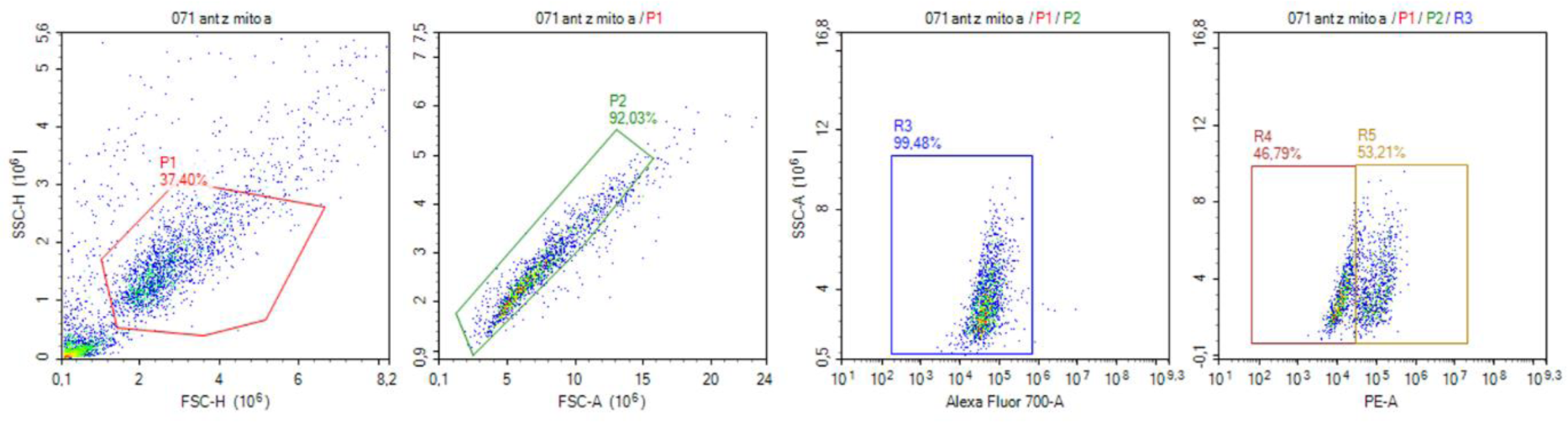
Gating strategy for response to H₂O₂, tBHP and antimycin A challenges. Five thousand events were counted and gated for SSC vs FSC, later exclusion of doublets and viability by ZombieNIR (Alexa Fluor 700).

