## Supplementary for "Disrupted stemness and redox homeostasis in mesenchymal stem cells of neonates from mothers with obesity: implications for increased adiposity"

### SUPPLEMENTARY MATERIAL

**Supplementary Table S1. Primers sequences and amplification conditions for RT-qPCR**

| mRNA Target | Forward Sequence | Reverse Sequence | Efficiency (%) |
|---|---|---|---|
| *SOD1* | GGTGTGGCCGATGTGTCTAT | CCTTTGCCCAAGTCATCTGC | 93 |
| *SOD2* | TGGGGTTGGCTTGGTTTCAA | TAGTAAGCGTGCTCCCACAC | 93 |
| *GPX1* | AGTGCGAGGTGAACGGTGCG | GGGGTCGGTCATAAGCGCGG | 94 |
| *CAT* | GTGCGGAGATTCAACACTGCCA | CGGCAATGTTCTCACACAGACG | 100 |
| *RPLP0* | AATCTCCAGGGGCACCATTG | GAACACCTGCTGGATGACCA | 100 |
| *GAPDH* | AGCCAACATCGTCAGACAC | GCCCAATACGACCAAATCC | 100 |
| *B2M* | GGTTTCATCCATCCATCCGACATT | ACGGCAGGCATACTCATCTT | 100 |

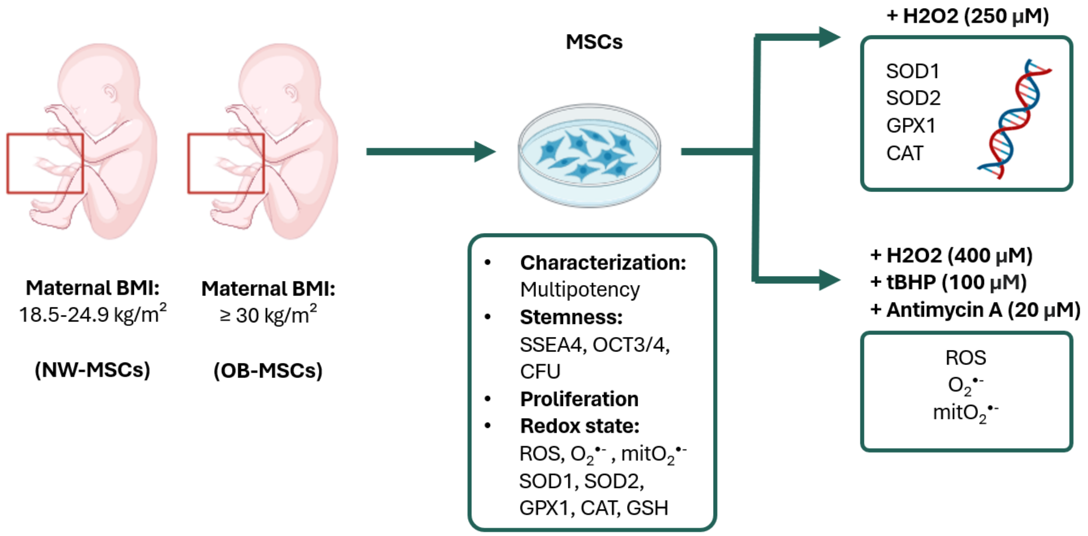


**Supplementary Figure S1. Study design.** MSCs were isolated from the Wharton's jelly of umbilical cords obtained from neonates of mothers with normal weight (NW-MSCs=15) and mothers with obesity (OB-MSCs=15). These MSCs were cultured characterized and further analysed to assess their stemness potential: SSEA4, OCT3/4, clonogenicity; and proliferation capacity. Further, we assessed redox parameters: reactive oxygen species (ROS), superoxide ($O_2^{\bullet-}$), mitochondrial superoxide (mit$O_2^{\bullet-}$) levels, gene expression of antioxidant enzymes and glutathione. Additionally, MSCs were subjected to pro-oxidative challenges: 250 μM $H_2O_2$ was used to induce changes in the gene expression of antioxidant enzymes, while 400 μM $H_2O_2$, 100 μM tert-butyl hydroperoxide (tBHP) or 20 μM Antimycin A, were used to induce intracellular ROS or $O_2^{\bullet-}$.

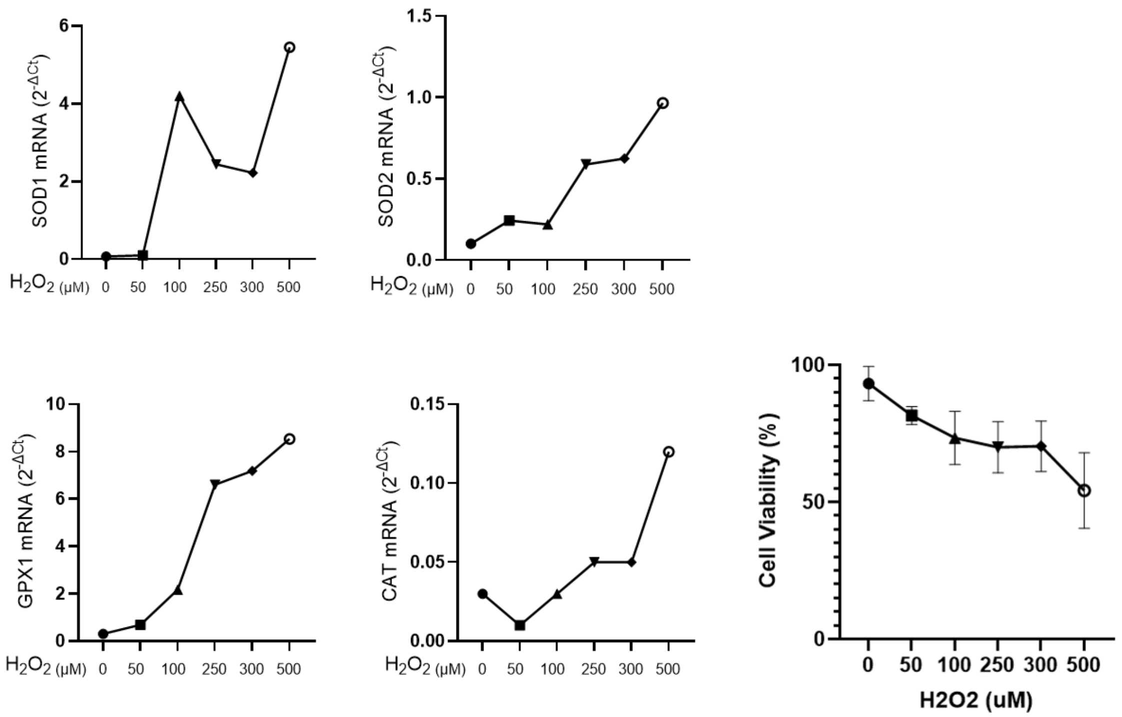


**Supplementary Figure S2. $H_2O_2$ challenge and viability assays in NW-MSCs.** Concentration-response to $H_2O_2$ experiments in NW-MSCs for standardization of an oxidative challenge. Left: gene expression of SOD1, SOD2, GPX1 and CAT was assessed after six hours of incubation with $H_2O_2$. Right: cell viability was evaluated with trypan blue in response to $H_2O_2$.

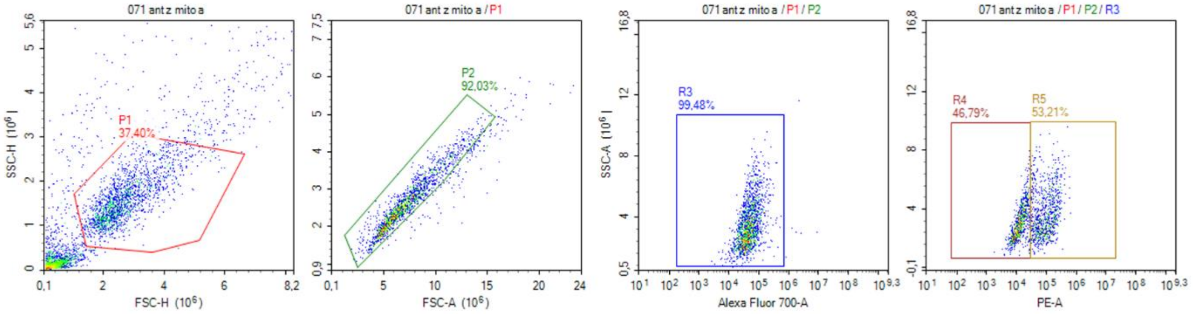


**Supplementary Figure S3: Gating strategy for response to $H_2O_2$, tBHP and antimycin A challenges.** Five thousand events were counted and gated for SSC vs FSC, later exclusion of doublets and viability by ZombieNIR (Alexa Fluor 700).
